# Genetic variability in the promoter region of TNF-α gene in the reservoir of Junin virus, Calomys musculinus (Rodentia, Cricetidae): Short communication

**DOI:** 10.1101/113688

**Authors:** Marina B. Chiappero, Imanol Cabaña, Gladys E. Calderón, Cristina N. Gardenal

**Author notes:** Corresponding author: Marina B. Chiappero.

## Abstract

In this study, we assessed the genetic variability of the promoter region of TNF-α gene in three natural populations of the cricetid rodent *Calomys musculinus*. This species is the natural reservoir of Junin virus, the etiological agent of Argentine Hemorrhagic fever. We found different levels of variability and varying signatures of natural selection in populations with different epidemiological histories.

---

In the last decades, South America has experienced the emergence of several zoonotic diseases produced by ARN viruses of the families Arenaviridae and Bunyaviridae (Enría and Pinhiero, 2000). The first of these diseases to be identified and studied was Argentine hemorrhagic fever (AHF), reported since the decade of 1950 in central eastern Argentina. Its etiological agent is the arenavirus Junin (JUNV) and the natural reservoir is the cricetid rodent *Calomys musculinus* (Parodi et al., 1958; Sabattini and Contigiani, 1982).Transmission of JUNV among rodents is mainly horizontal (Mills et al., 1992). When infected, *C. musculinus* individuals experience an acute phase characterized by active virus replication. Thereafter, some individuals clear the infection while others shift to a persistent (chronic) phase, remain infected for life and continuously shed the virus into the environment through body fluids, infecting other rodents and man. Chronically infected rodents are asymptomatic and are the primary responsibles for the maintenance of JUNV in nature (Enría et al. 2006).

The endemic area of AHF (defined by the occurrence of the disease in humans) has been expanding gradually since the discovery of the disease and continues to expand (Enría et al., 2002; XVIII Annual Meeting of the National Programme of Control of the Argentine Hemorrhagic Fever, 2015; http://www.anlis.gov.ar/inevh/?pageid=215). Three epidemiological zones are recognized, according to disease incidence: a non-endemic zone with no cases, an epidemic zone characterized by the continuous presence of human cases with a mean incidence greater than 2.0/10000 inhabitants, and a historic zone with a mean incidence less than 2.0/10000 inhabitants (Mills et al., 1991). A given area usually goes from non endemic (no viral presence), to epidemic for 5 to 10 years, and thereafter infection gradually declines to only sporadic cases and the locality is considered historic (Calderón 2004; García et al., 2000).The influence of different factors have been studied to explain this phenomenon: virulence variability of JUNV circulating strains (Calderón, 2004), the population dynamics of the rodent (Polop et al., 2007; Porcasi et al., 2005; among others), and patterns of migration among reservoir populations (Chiappero et al., 2003, 2011; González Ittig et al., 2004).However, an important aspect has remained untreated in the species. Genes of the immune system have been studied in natural populations of numerous animal species, to provide insights into the relative influence of their genetic variation on the interactions host-pathogen (Deter et al. 2008; Guivier et al., 2014). Tumor Necrosis Factor-alpha (TNF-α) is a cytokine, produced by activated macrophages, involved in the resistance against certain infectious agents including viruses. Guivier et al. (2010b) found geographic variation in the distribution of 16 SNPs in the promoter region of TNF-α gene in the Puumala virus rodent reservoir in Europe, *Myodes glareolus.* Two genotypes, significantly associated to increased gene expression, were more frequent in non endemic areas. Unfortunately, the precise mechanisms of development of chronic infection in *C*. *musculinus* by JUNV are poorly known, partly due to the logistic and biological difficulties of working with these models (Golden et al., 2015), but TNF-α gene may play a role. Heller et al. (1992) found that TNF-α levels were markedly high in human FHA patients, particularly in fatal cases. Coulombié et al. (1986) found that macrophages are directly related to the resistance of adult rodents to JUNV infection. Coto et al. (1993) determined that peritoneal macrophages were involved as a source of virus serological variants in vivo, able to avoid immune elimination and to reach the brain, the major target organ of JUNV persistent infection. Therefore, this gene may be implicated in the course of infection both in humans and in the rodent reservoir. Here, we assessed intra and interpopulation genetic variability of the promoter region of TNF-α gene in three natural populations of *C. musculinus* to explore its possible link with their epidemiological history.

Total genomic DNA was extracted from 30 *C. musculinus* from 3 natural populations with different epidemiological histories (Fig. 1): Oliveros (non-endemic; N = 7), San Pedro (epidemic; N = 11) and Pergamino (historic; N = 12), using a standard salt purification procedure followed by ethanol precipitation (protocol 1 of Bruford et al. 1992). The promoter region of TNF-α gene was amplified by the polymerase chain reaction (PCR) using specific primers designed from conserved regions between *Rattus norvegicus* (D00475.1) and *Mus musculus* (M20155.1): Mur-TNFp-F: 5'-CCAGGGCTGAGTTCATTCCCTCTGG-3' and Mur-TNFp-R: 5'-CGGATCATGCTTTCYGTGCTCATGGTGTC-3'. Polymerase chain reactions were carried out in 50 μl final volume, containing 1 unit of Taq polymerase (Biolase, Bioline USA Inc.), 5ng of genomic DNA, 2.5 mM MgCl_2_, 0.2μM of each primer, and 0.2 mM of each dNTP (GE Healthcare, Uppsala, Sweden). The cycling program consisted of an initial denaturation step at 94° C for 5min, followed by 30 cycles of denaturation at 92° C for 1 min, 1 min at 58°, and extension at 72° C for 1 min; the final extension step consisted of 5 min at 72° C. PCR products were purified and sequenced in both directions at Macrogen (Seoul, Korea). Sequences were verified and corrected by eye using Chromas Lite 2.01 (http://www.technelvsium.com.au). aligned using the ClustalW algorithm implemented in Mega 6 (Tamura et al. 2013) and trimmed to equal lengths where necessary. Identity of fragments was confirmed using the BLAST search program of the National Centre for Biotechnology Information website (http://www.ncbi.nlm.nih.gov).For sequences containing heterozygous peaks, the allelic phase of each haplotype was resolved using the PHASE algorithm implemented in Mega 6.

**Fig.1.**
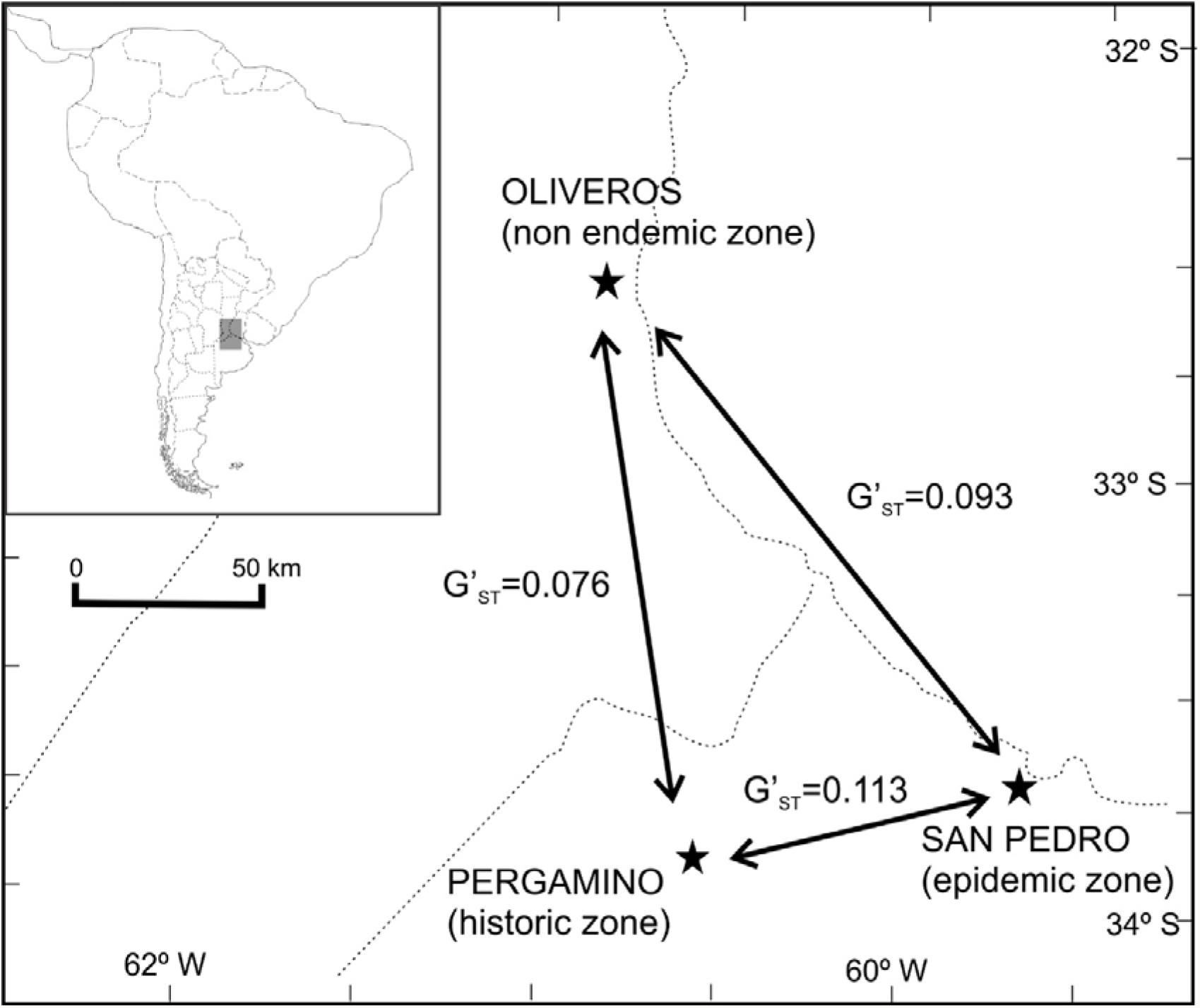
Geographic location and genetic differentiation of populations analized. Genetic differentiation was calculated as G'_ST_ (Hedrick 2005), most adequate for loci with high levels of heterozygosity. Significance of G'_ST_ values was assessed through 10000 permutation. Calculations were performed with Genalex 6.51 (Peakall and Smouse 2012).All G'ST values were statistically significant at the 0.001 level.

A fragment of 624 pb was analyzed in all individuals, containing 11 polymorphic sites that defined 15 alleles. Sequences for all alleles were submitted to GenBank (accession numbers KY679018 to KY679032). Identity of the alleles was confirmed using the BLAST search program of the National Centre for Biotechnology Information website (http://www.ncbi.nlm.nih.gov), as all showed the highest similarity with the 5′ end of the published sequence of TNF-α gene of *Peromyscus leucopus* (58% coverage, 92% identity; M59233.1) and *Mus musculus* (99% coverage, 81% identity; GQ917239.1, U68414.1).

In all populations, genotypic frequencies in this locus were in Hardy-Weinberg equilibrium. Only 3 alleles were shared by all populations; most were exclusive of a population (Table 1). These calculations were performed with Arlequin 3.5 (Excoffier and Lischer 2010). The three populations were significantly differentiated at this locus (Fig. 1). The population of Pergamino (historic zone of AHF) showed the lowest values of h_e_ and a_r_, while the epidemic and non endemic populations showed similar levels of genetic variability (Table 2).

**Tabel 1:**
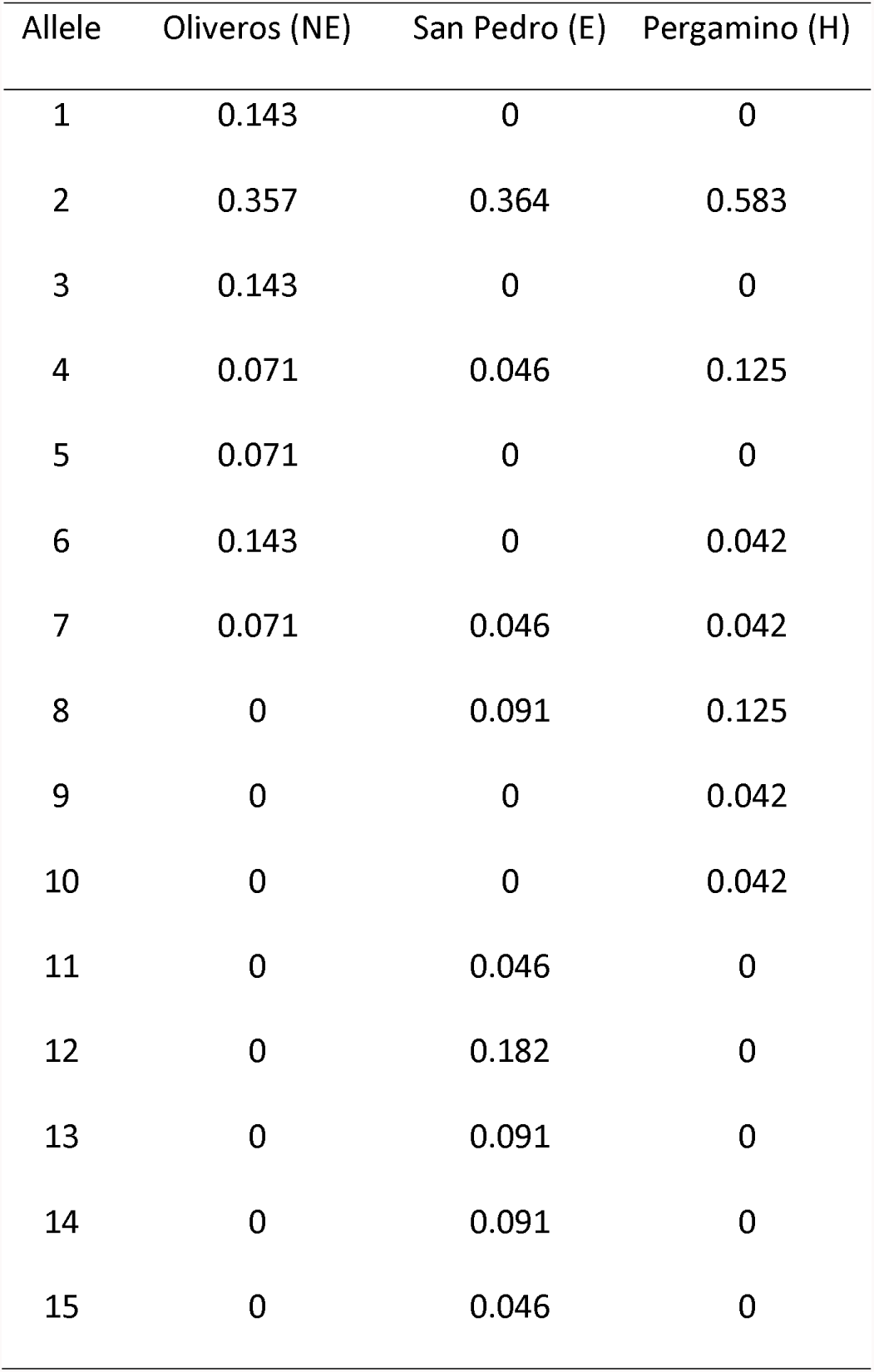
Relative allelic frequencies of TNF-α promoter gene in three natural populations of *C. musculinus* with different epidemiological histories. NE: non endemic zone; San Pedro: epidemic zone; H: historic zone.

**Tabel 2:**
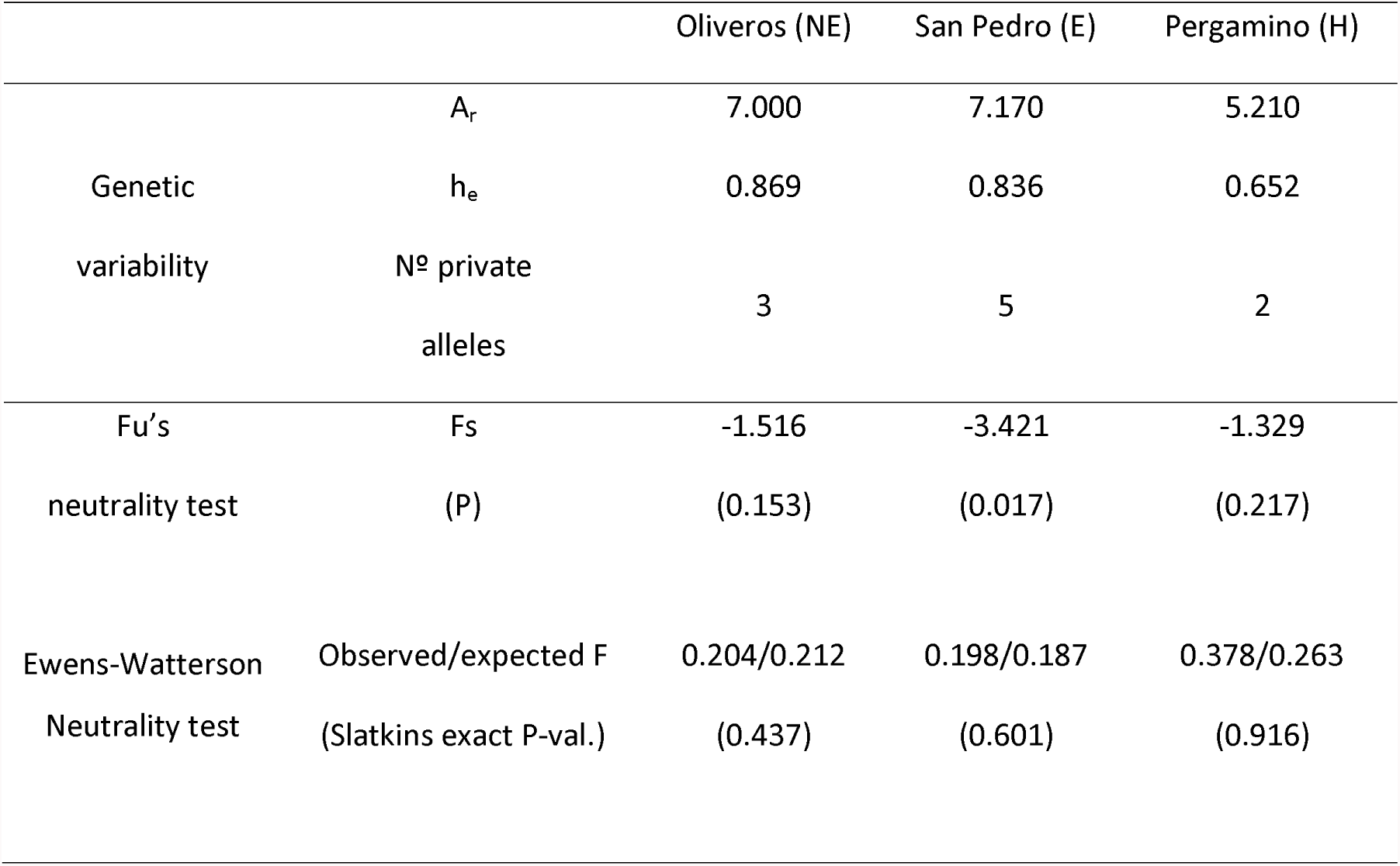
Genetic variability and tests of selective neutrality in three populations of C. musculinus. A_r_: allelic richness, calculated with FSTAT (Goudet, 2001); he (expected heterozygosity) and number of private alleles were calculated with Genalex 6.5 (Peakall and Smouse, 2012). Fu's and Ewens-Waterson neutrality tests were calculated with Arlequin 3.5 (Excoffier and Lischer, 2010).

Two neutrality tests were performed to examine if natural selection is acting on this gene in *C. musculinus* populations in two different time frames (Turner et al. 2012). Fu′s (1997) Fs test, examines evidence for selection acting over the long term, by comparing the observed number of haplotypes with the expected number of haplotypes under a neutral coalescent model with the same θ _π_. Ewens-Watterson test (Ewens 1972;Watterson 1978), on the other side, examines evidence of contemporary selection, by comparing the observed homozygosity in the sample against its expected distribution under neutrality, for a given number of alleles and population size. Significance of both tests was assessed using 10000 permutations in Arlequin 3.5. The three populations showed negative values of Fs, significant only for the epidemic population (Table 2). A negative value of Fs is evidence of an excess number of alleles, as would be expected from genetic hitchhiking. Ewens-Watterson tests, on the other side, yielded non significant results (Table 2), but the historic population showed a marginally non significant excess of homozygotes. An increase in homozygotes compared to that expected under neutral evolution may be due to positive or negative selection. However, it has to be kept in mind that these tests rely on the assumption that the populations are in mutation drift equilibrium. Populations that experienced a recent demographic expansion would, for example, show also negative Fs values. González Ittig and Gardenal (2004) and González Ittig et al. (2007) found evidence of a recent population expansion in *C. musculinus* from central Argentina using two mitochondrial genes as genetic markers. Unfortunately, they did not calculate Fs for each population separately, perhaps due to the small sample sizes used in the study. However, if the negative Fs in the epidemic population were due to a population expansion, this signature would be present in all populations, and with other kind of genetic markers such as neutral microsatellite loci(Beaumont and Balding 2004). Chiappero et al. (unpublished results) estimated effective population sizes in 14 populations of *C. musculinus* with different epidemiological histories using 6 microsatellite loci. They found that effective population sizes were smaller in historic populations, compared to epidemic and non endemic populations. Therefore, the negative value of Fs obtained here is probably not the result of a population expansion but of a long term selective pressure.

Chiappero et al. (unpublished results) suggested that the lower reproductive success of females with chronic infection, compared to those that develop an acute infection, would lead to an increment of the proportion of the latter expenses of that of the first kind. Since the persistence of the virus in a rodent population depends on the chronically infected rodents, virus would slowly clear from the population, helping to explain the changing incidence of the disease. While the preliminary results of the present work are promising, we need to increase sample sizes and number of populations analyzed, and to screen other genes, potentially involved in the immunological response to arenaviruses like other cytokines, interferons, Toll-like receptors and MHC class II genes (Guivier et al. 2010a; Koma et al., 2013; Turner et al., 2011, 2012).

## Competing interests

The authors declare that they have no competing interests.

## Acknowledgements

This study was supported by a grant from FONCyT (PICT-2012 N 1275). The authors gratefully thank Luciana Plum for her assistance in laboratory work.

